# Milieu-Initiated Inversion of the Aqueous Polyproline II/Beta Propensity in the Alanine Tripeptide: Energetics Origin of the Onset of Amyloid Formation

**DOI:** 10.1101/238006

**Authors:** Noemi G. Mirkin, Samuel Krimm

**Affiliations:** LSA Biophysics, University of Michigan, 930 N. University Ave., Ann Arbor, MI 48109

## Abstract

Extending our earlier analogous study of the alanine dipeptide (ADP), we have now analyzed the effect of the external environment on the polyproline II (P) and β relative energies, the P/β propensity, of the alanine tripeptide (ATP). Ab initio calculations of ATP(H_2_O)_19_ and ATP(H_2_O)_19_(HCl) exhibit the same propensity inversion as in ADP: in the pure-water case the PP conformation is favored while the addition of the HCl molecule results in the ββ conformation being of lower energy. A comparison, following an intermediate insertion and departure of an HCl molecule, shows that the energy of a hydrogen-bonded (H_2_O)_19_βATP::βATP(H_2_O)_19_ structure is lower than that of the sum of two separate PP systems, i.e., that the aggregated state of the peptide is favored. This arises from the basic physical response to their total environmental influences. Questions about quantitative results from molecular dynamics simulations, obviously needed to analyze longer chains and other side chains, are addressed via rigid-water calculations. The desirability of basing studies of amyloid formation on our proposed alternative milieufolding paradigm is discussed.

## INTRODUCTION

It is now generally accepted, as proposed^1^ and substantiated^2^, that a significant contribution to the circular dichroism (CD) spectra of unordered polypeptides and proteins in aqueous solution (a strong negative band near 200 nm and a weak positive band near 220 nm) comes from locally ordered chain conformations close to that of the three-fold left-handed helix of polyproline II (P). Our ab initio studies of a hydrated alanine dipeptide (ADP) indicated that this preference over that of the nearby β conformation is determined primarily by the difference in the interaction energies associated with their peptide-water and water-water hydrogen-bonded structures^3^. The idea that the relative P and β energies (i.e. P/β propensity) depend on the specific features of these structures motivated a detailed study of the possible influence on this propensity of other species in the aqueous environment and resulted in our analogous investigation of this property of hydrated ADP with an added HCl molecule. Our finding was that the P/β propensity becomes inverted, that is the energy of the β conformer is now lower than that of the P conformer^4^.

The recognition that this could have implications for the formation of plaques in amyloidosis diseases was further supported by showing that a hydrated hydrogen-bonded two-β-peptide state is energetically favored over one in which the two initial hydrated P components are separate, an indication that aggregation of β chains is to be expected in favorable environments. Although this emphasized the importance of accepting a peptide-medium paradigm as the basis of structure determination, it was also clear that further detailed study of peptides was needed. Aside from the likelihood that other environmental conditions besides pH could initiate such changes, it would be necessary to explore the influence of factors such as chain length and side-chain composition. As a start in this direction, we report here on an ab initio study of the P/β propensity in the hydrated alanine tripeptide (ATP) in the absence and presence of an HCl molecule. We also consider the important issue of reproducing by molecular dynamics the accuracy of such basic quantum-mechanical calculations in needed studies of longer chains and other side chains, and deal as well with the implications of our results for achieving a deeper understanding of amyloid formation.

## CALCULATIONAL METHODS

The ATP molecule, CH_3_-CONH-C_1_H(CH_3_)-CONH-C_2_H(CH_3_)-CONH-CH_3_, hydrated with water was minimized in the following way: 1) the peptide was constructed with the canonical values of φ,ψ (P: −75°, 145° and β: −134°, 145°); 2) first-layer waters were added to the ubiquitous peptide hydrogen-bonding positions, i.e., two to the CO and one to the NH, comprising a total of 9 water molecules, and the entire system was minimized; 3) using the new φ,ψ as a starting point and adding sufficient waters to complete the second layer, 10 in this case for a total of 19, the system was again totally relaxed and minimized. As before^4^, the global and some local minima of each conformation were determined. For the ATP(H_2_O)_19_(HCl) system, as previously^4^, the HCl was added to the above minimized structures in a variety of ways: as an intact molecule initially located in numerous positions with respect to the water layers and as initially separate ions in various locations and the total system was minimized. In all cases the final structures had separated ions imbedded in the water environment.

Calculations were done with Gaussian 09^5^, with the dispersion-corrected wB97X-D functional^6^ and the 6-31++G** basis set. In addition, a Polarized Continuum Model (PCM) reaction field calculation was done on the final minima in order to assess the impact of general polarization contributions from the surrounding medium on the final structures and energies. Atomic charges were determined as Natural Population Analysis (NPA) charges.

## RESULTS AND DISCUSSION

### Alanine Tripeptides

The energy difference of importance in the ATP systems is that between the P_1_P_2_ and β_1_β_2_ conformations, i.e., ΔE = E(P_1_P_2_) – E(β_1_β_2_). The P/β propensity inversion is defined by the sign change of ΔE. This quantity for ATP(¾O)i9 is shown in Table 1. The – 1.80 kcal/mol for β_1_β_2_ indicates that its energy is higher than that of P_1_P_2_ (remembering that each of the individual energies is negative). This value is in line with the result of −1.72 kcal/mol for the comparable ΔE of ADP(H_2_O)_12_^4^ in indicating the preferred stability of the P_1_P_2_ conformation in an all-aqueous environment (even though the energy of an isolated β_1_β_2_ structure with the φ,ψ of the hydrated structure is 1.28 kcal/mol lower than that of the comparable P_1_P_2_). Such an energy difference is associated with the different total interaction energies (i.e., peptide-water and water-water) in the two structures^4^, part of which shows up as a difference in total charge on the peptide, in this case −2 me on P_1_P_2_ and 37 me on β_1_β_2_ (compared to zero for the neutral isolated structures). The inclusion of a PCM reaction field in the calculation does not change the φ,ψ appreciably but the ΔE increases to −2.62 kcal/mol and, presumably as a result of the additional effects of polarization and charge transfer through water-peptide hydrogen bonds^7, 8^, the relevant peptide charges become more positive, at 28 me and 59 me, respectively.

**Table 1.** Properties of ATP(H_2_O)_19_ and ATP(H_2_O)_19_(HCl)

| System | $(\phi_1, \psi_1)$ | $(\phi_2, \psi_2)$ | $\Delta E^a$ | $q(\text{pep})^b$ | $q(\text{Cl}^-)^b$ |
| --- | --- | --- | --- | --- | --- |
| $\text{ATP}(\text{H}_2\text{O})_{19}$ | | | | | |
| $\text{P}_1\text{P}_2$ | $(-81^\circ, 135^\circ)$ | $(-65^\circ, 146^\circ)$ | 0 | -2 | |
| $\beta_1\beta_2$ | $(-158^\circ, 150^\circ)$ | $(-163^\circ, 128^\circ)$ | -1.80 | 37 | |
| $\text{ATP}(\text{H}_2\text{O})_{19}$ PCM | | | | | |
| $\text{P}_1\text{P}_2$ | $(-83^\circ, 138^\circ)$ | $(-67^\circ, 141^\circ)$ | 0 | 28 | |
| $\beta_1\beta_2$ | $(-154^\circ, 140^\circ)$ | $(-153^\circ, 121^\circ)$ | -2.62 | 59 | |
| $\text{ATP}(\text{H}_2\text{O})_{19} (\text{HCl})$ | | | | | |
| $\text{P}_1\text{P}_2$ | $(-92^\circ, 115^\circ)$ | $(-75^\circ, 137^\circ)$ | 0 | -30 | -806 |
| $\beta_1\beta_2$ | $(-147^\circ, 148^\circ)$ | $(-146^\circ, 116^\circ)$ | 2.47 | -6 | -814 |
| $\text{ATP}(\text{H}_2\text{O})_{19} (\text{HCl})$<br>PCM | | | | | |
| $\text{P}_1\text{P}_2$ | $(-91^\circ, 118^\circ)$ | $(-74^\circ, 132^\circ)$ | 0 | -10 | -837 |
| $\beta_1\beta_2$ | $(-146^\circ, 144^\circ)$ | $(-144^\circ, 120^\circ)$ | 2.86 | 36 | -862 |

<sup>a</sup> $\Delta E = E(\text{PP}) - E(\beta\beta)$ , kcal/mol.<sup>b</sup> NPA charge in me.

The situation for ATP(H_2_O)_19_(HCl), Table 1, is significantly different from the ATP(H_2_O)_19_ case: as with ADP(H_2_O)_12_(HCl), the P/β propensity is inverted, with the β_1_β_2_ conformer now being 2.47 kcal/mol more stable than P_1_P_2_. The peptide charges are significantly more negative, at −30 me and −6 me respectively, but again the β_1_β_2_ charge is more positive than that on P_1_P_2_. The inclusion of a PCM reaction field results in a modest increase in ΔE, to 2.86 kcal/mol, and the peptide charges again become more positive. It is interesting that the positive charge increase from P_1_P_2_ to β_1_β_2_ is accompanied (although not quantitatively) by an increase in the negative charge on the Cl^-^ ion.

The inversion of the P/β propensity in the HCl case was found to favor peptide aggregation in ADP(H_2_O)_12_^4^ (an inadvertent reversal of the P and β values occurred in the 2.48 and 0 of Table II in reference 4), and we examined if the same is true for the tripeptide. The result can be inferred from the following energy comparison. We focus on two individual ATP molecules: 1) in a pure aqueous environment both ATPs are preferably in the P_1_P_2_ conformation, i.e., “solubilized”; 2) an HCl molecule attaches to one ATP, which then assumes the more favorable β_1_β_2_ conformation; to reach the final state we now suppose that the other P_1_P_2_ peptide converts to ββ and it hydrogen-bonds (:::) to the β_1_β_2_ peptide to form the ATP:::ATP(H_2_O)_38_(HCl) complex, which will lower the energy^4^; 3) the HCl molecule departs and we compare the total energy of this system with that of 1), the originally separated P_1_P_2_ components. The energy of 3) was determined by forming the optimally hydrogen-bonded antiparallel ATP:::PTA pair (comparable parallel-chain pairs are also likely for longer chains), adding the necessary water molecules in the initial layers, relaxing the system and minimizing, followed by subsequent additional water layers with relaxation and minimization until a final minimized (H_2_O)_19_(ATP:::PTA(H_2_O)_19_ structure was achieved. The energy of this structure is 22 kcal/mol lower than that of 1) (a PCM calculation maintains the sign of this difference), which is not unexpected since there are more non-hydrogen-bonded water OH groups in 1) than in 3) that are not compensated by the few peptide-peptide hydrogen bonds in 3) or by the small energy cost of converting both P_1_P_2_ to β_1_β_2_. In the longer chain cases we expect that similar prior constraints on near-peptide water molecules in the P conformation are released in the β conformation thus leaving them to participate in stronger water-water hydrogen-bonding arrangements. In effect, the HCl has acted like a chaperone-type enabler to the transition from a solubilized system to one in which the peptide components tend to aggregate. If these structures precipitate, removing the peptides from circulation, the system will re-equilibrate and be repeatedly driven toward aggregation, a situation that is extremely unlikely dynamically in 1). Of course, if both molecules in 1) have HCl entrained and therefore in ββ conformations then the probability of forming an aggregate is enhanced. (The true propensities are obviously determined by the free energies, but if the energy profiles in the ATP cases are similarly shaped, as in the ADP case^4^, implying that the entropies of the two structures are about the same, the free energies will track the enthalpies.) It might be thought that this result is just an abstract theoretical exercise, but there has long been strong x-ray diffraction evidence of its occurrence in low molecular weight synthetic polypeptides, with the formation of cross-beta structures^9^ (in which the local peptide chain axis is perpendicular to the global fibril axis), and indeed is a long-known property of proteins^10, 11^. It thus has to be concluded that such aggregation is a natural consequence of originally solubilized peptide conformations in the presence of appropriate milieus.

As noted before^4^, the magnitude of the calculated aqueous P/β propensity inversion could depend on the chain length and the nature of the side chain. Since these properties need to be studied, and obtaining such correlations from the ab initio methods used here is clearly not a viable computational procedure, the necessary route has to be through molecular dynamics (MD) calculations (which in addition can also provide free energies). Given the sensitive dependence of the results on the detailed nature of the various hydrogen-bonding interactions in such systems, it is appropriate to ask if current MD force fields are up to this task, even qualitatively at the level of ab initio calculations. Since charge transfer is not incorporated in present force fields (and, as shown in Table 1, the relative magnitudes could be significant) and even charge flux terms (championed by some to improve accuracy^12, 13^) remain to be dealt with in current force fields, this is an open question. A simple example of the problem is seen from the results of calculations with rigid water molecules, a common assumption in many present MD simulations^14^. For ATP(H_2_O)_19_, although the φ,ψ are almost identical to the flexible water case, ΔE = – 3.86 kcal/mol for the rigid water case compared to the –1.80 kcal/mol for flexible water (with PCM, again with almost identical φ,ψ, the comparable values are −6.55 kcal/mol and −2.62 kcal/mol respectively). The peptide charges are also significantly different: −17 me for the P_1_P_2_ and 21 me for the β_1_β_2_ conformations compared to −2 me and 37 me respectively for the flexible water case (with PCM the values are 7 me for P_1_P_2_ and 38 me for β_1_β_2_ in the case of rigid waters compared to 28 me and 59 me respectively in the case of flexible waters.) This level of sensitivity to local interactions highlights the need to develop “micro-fine-grained” force fields^4^ to achieve reliable descriptions of P/β propensity from MD simulations. Structure agreement itself is a necessary but is not a sufficient requirement to guarantee energy accuracy.

### Amyloid Formation

There are almost 70 human diseases, both neurodegenerative (such as Alzheimer’s and Parkinson’s) and non-neuropathic (such as type II diabetes), that are now associated with the formation of insoluble amyloid fibrils^15^. Structural studies by many physical techniques^16^ find that these fibrils, and the many types of aggregates (polymorphs) that are formed from them, are based on a local cross-β chain conformation, said to result from the “misfolding” of their peptides. In the case of the Alzheimer’s Aβ peptides, these are of about 40 residues, derived from enzymatic cleavage of its amyloid precursor protein, and are biologically functional in their normally soluble state^17^. The formation of the insoluble amyloid plaque is known to be basically independent of the amino acid sequence and is accepted as an inherent property of the polypeptide chain^15^ (it even being speculated that the cross-β motif might have been a common ancestor of protein folds^18^). Much has been learned about factors influencing amyloid properties, including those such as hydration defects in the cerebrospinal fluid, which can interfere with the normal clearance of Aβ from the brain^19, 20, 21^. Ions such as Zn^2+ 22^ and Al^3+ 23^ enhance amyloid formation while many molecules are found to inhibit the formation of Aβ (H_2_S^24^, vitamin B12^25^, and ascorbic acid^26^ among others). However, the basic and still unanswered question has been what physical property can lead to the formation of an ordered peptide structure from a supposedly unordered chain state. Our studies indicate that the fundamental answer lies in the dependence of the P/β propensity in the chain on the specific nature of its environment.

To further evaluate this view, it is important to first understand the detailed structural nature of the initial soluble state. Such peptide chains have generally been described as being in an intrinsically disordered^15^ or random state, especially from CD spectral analyses. Unfortunately, these have been based on an early assignment to the random state^27^ of a CD spectrum that is now recognized as significantly associated with the PPII conformation^1, 2^. A CD study of the Aβ(12-28) fragment^28^ makes this very clear through the temperature dependence of its spectrum (enhanced PPII contribution at lower temperature) and the presence of an isodichroic point^29^ between the PPII spectrum and one consistent with a basically unordered chain^30^. This composite nature of such soluble systems is further supported by CD studies of Aβ(1-40)^31^. Since PPII conformations in adjacent chains cannot effectively hydrogen-bond to each other, their presence in the peptide chain is clearly antagonistic to chain aggregation and thus to the formation of cross-β fibrils.

The possibility of amyloid formation then follows from two important results of our ADP and ATP studies on the influence of other species in the aqueous environment. First, since the P/β propensity favors the PPII conformation in a totally aqueous environment, which is obviously not the case in a normal brain, it must be true that certain compositional features of this natural environment still permit this propensity to prevail. Second, since we find that some added species (or even changes in pH^32^) alter the propensity in favor of the β conformation and thus can lead to chain aggregation, it is likely that such an addition to the milieu (including seeding by externally introduced amyloid^33^) constitutes the initiation of amyloid onset. A shift from the present misfolding to a more useful milieufolding paradigm is therefore an important conceptual step in developing a basic understanding of amyloid development.

## CONCLUSIONS

Our previous ab initio studies on the hydrated alanine dipeptide^4^ and our present similar analysis of the comparable alanine tripeptide show that the relative energies of the PPII and β conformations, the P/β propensity, depends on the local environment, favoring the P and P_1_P_2_ conformers in the ADP and ATP cases, respectively, in a pure aqueous environment and the β and β_1_β_2_ conformers in the case of an added HCl molecule in the system. We believe that other molecules can have the same effect. Since our ab initio approach cannot be used to establish the magnitude of this dependence for longer chains and other side chains, as is desirable, such studies will necessarily have to depend on molecular dynamics simulations. It is quite possible that present force fields are not able to provide a quantitatively reliable description, and we show that this is indeed a problem in the case of an ab initio calculation of ATP with rigid rather than flexible waters. Force fields will need to demonstrate their ability to reproduce such obviously subtle hydrogen-bond-dependent effects.

Implementation of our proposed alternative paradigm of protein milieufolding diseases can lead to better-targeted experimental approaches in studying the onset phase of amyloid formation. We have pointed to the desirability, since neurodegenerative diseases are usually age-related, of monitoring compositional changes in the cerebrospinal fluid that, through lifestyle and/or environmental factors, influence P/β propensity^4^. It may also be the case that blood-brain barrier (BBB) changes, already known to be associated with Alzheimer’s^34^, are involved, perhaps through gut microbiota influence on BBB permeability^35^. Detailed knowledge of the initial physical steps in favoring β over PPII polypeptide chain conformations most likely holds the keys to enabling effective interventions in the formation of biologically destructive amyloid deposits.

## ACKNOWLEDGEMENT

A major portion of our calculations was made possible by the donation of time on the computational facilities of Charles L. Brooks III. For this we are deeply grateful.

